# Loss of mitochondrial Chchd10 or Chchd2 in zebrafish leads to an ALS-like phenotype and Complex I deficiency independent of the mt-ISR

**DOI:** 10.1101/2022.05.02.488746

**Authors:** Virginie Petel Légaré, Christian J. Rampal, Mari J. Aaltonen, Alexandre Janer, Lorne Zinman, Eric A. Shoubridge, Gary A.B. Armstrong

## Abstract

Mutations in *CHCHD10* and *CHCHD2*, coding for two paralogous mitochondrial proteins, have been identified in amyotrophic lateral sclerosis (ALS), frontotemporal lobar degeneration (FTD), and Parkinson’s disease (PD). Here we investigated the biological roles of these proteins during vertebrate development using knockout (KO) models in zebrafish. We demonstrate that loss of either or both proteins leads to a motor impairment, reduced survival, and compromised neuromuscular junction (NMJ) integrity in larval zebrafish. Compensation by Chchd10 was observed in the *chchd2*^-/-^ model, but not by Chchd2 in the *chchd10 ^-/-^* model. The assembly of mitochondrial respiratory chain Complex I was impaired in *chchd10 ^-/-^* and *chchd2 ^-/-^* zebrafish larvae, but unexpectedly not in the double *chchd10 ^-/-^* & *chchd2 ^-/-^* model, suggesting that reduced mitochondrial Complex I cannot be solely responsible for the observed phenotypes, which are generally more severe in the double KO. Activation of the mitochondrial integrated stress response (mt-ISR) was only observed in the double KO model, possibly implicating this pathway in the recovery of the Complex I defect, and suggesting that Complex I assembly defect in our single KO is independent of the mt-ISR. Our results demonstrate that both proteins are required for normal vertebrate development, but their precise molecular function in the mitochondrial biology of motor neurons remains to be discovered.

## Introduction

ALS is a devastating neurodegenerative disease characterized by the progressive degeneration of lower and upper motor neurons in the spinal cord and motor cortex, that in some cases extends to the frontal and temporal cortexes where it manifests as FTD (1). While most forms of the disease occur sporadically (sALS), a small subset of patients (∼10%) have a familial history (fALS). Among the many defects observed in ALS, mitochondrial abnormalities have long been documented as pathological features, with patients presenting with mitochondrial structural defects, altered mitochondrial respiration, and increased oxidative stress (2–4). Yet, whether mitochondrial dysfunction plays a central role in the disease or arises as a downstream consequence of other cellular abnormalities remains to be fully elucidated. In 2014, the first dominantly inherited variants of the nuclear encoded mitochondrial protein coiled-coil-helix-coiled-coil-helix domain containing 10 (CHCHD10) were reported in a small proportion of both fALS, sALS and FTD cases (5) suggesting that mitochondrial protein abnormalities could play a direct role in the disease. Since these original findings, over twenty mutations in *CHCHD10* have been associated with a spectrum of diseases, including ALS, FTD, cerebellar ataxia and mitochondrial myopathy (5), Charcot–Marie–Tooth disease (6), PD (7), and Alzheimer’s disease (AD) (8).

In mitochondria, CHCHD10 localizes mostly near the cristae junctions of the mitochondrial intermembrane space (IMS) (5, 9), where it can form a complex of unknown function with its paralogue CHCHD2 (9, 10). CHCHD2 has also been linked to neurodegenerative diseases with rare autosomal dominant mutations in *CHCHD2* segregating with PD (11, 12), FTD (13) AD (13), and Lewy body dementia (14).

CHCHD2 and CHCHD10 share 58% sequence identity and have a common ancestor in yeast, suggesting that these proteins might have partially overlapping function (15). *In vitro* studies have reported that loss of either CHCHD2, CHCHD10 or both, impacts mitochondrial respiration (10, 16), survival/growth following exposure to various cellular stressors (9, 10, 16, 17), the mitochondrial integrated stress response (mt-ISR) (18–22), apoptosis (23, 24), as well as the transcription of the oxygen responsive gene *COX4I2* (25–27).

Conversely, *in vivo* studies employing mouse knockout (KO) models reported only minor defects following loss of either protein (19, 28, 29), though when aged, Chchd2 KO mice did develop Parkinsonian-like symptoms (30). More recently, a double KO mouse model was created, which presented with mitochondrial structural defects, cardiomyopathy, as well as activation of the mt-ISR (21). Other loss-of-function models, including hypomorphic and KO models of CHCHD10 and the CHCHD2 orthologue DmeI\CG5010 in *Drosophila melanogaster* (28, 31), a KO model of the CHCHD10 and CHCHD2 orthologue *har-1* in *Caenorhabditis elegans* (32), and a knockdown model (KD) of *chchd10* in *Danio rerio* (33), reported reduced survival and impaired motor activity, suggesting that mouse models might have a broader capacity to compensate for loss of CHCHD10 and/or CHCHD2. Animal model studies have largely focused on either protein in isolation or used models in which CHCHD10 and CHCHD2 are represented by only one orthologue leading to disparate conclusions concerning their biological function. The zebrafish model system offers the opportunity to manipulate both genes in isolation or in combination, as both *chchd2* and *chchd10* are well-conserved, with 65% and 67% identity to human, and 60% and 72% identity at the amino acid level, respectively. As zebrafish have already been successfully used to model mitochondrial pathology (34–38), we generated zebrafish CHCHD10 and CHCHD2 single and double KO models and explored the biological ramifications during development, enabling us to gain insight into their roles during vertebrate development.

## Materials and Methods

### Zebrafish Housing and Maintenance

Adult zebrafish (*D. rerio*) of the Tübingen long fin (TL) strain were maintained according to standard procedures (39) at 28.5 °C under a 14/10 light/dark cycle at the animal research facility of the Montreal Neurological Institute (MNI) at McGill University located in Montréal Québec, Canada. All experiments were performed in compliance with the guidelines of the Canadian Council for Animal Care and approved by the Animal Care Committee of the MNI.

### Cas9 mRNA and guide RNAs Synthesis

Synthesis of Cas9 mRNA and guide RNA (gRNA) was performed as previously described (40, 41). Briefly, a zebrafish codon optimized Cas9 (pT3TS-nCas9n, Addgene plasmid # 46757) was linearized with XbaI overnight and 1 μg of linear template DNA was used for *in vitro* transcription of mRNA using the T3 mMESSAGE mMACHINE® Kit (Invitrogen) followed by phenol-chloroform extraction and precipitation with ethanol. Selected gRNA target sites were identified using CRISPRscan (42) synthesized using the T7 MEGAscript kit (Invitrogen) and purified by phenol-chloroform extraction and ethanol precipitation. The following gRNA target sites were used (PAM site is underlined): Chchd10 gRNA target site: TGGCCGTTGGTTCAGCTGTT<u>GGG</u>; Chchd2 gRNA target site: <u>GGT</u>GACGACGTCCGCAACGACAG.

### CRISPR Mutagenesis and Screening of Founder Lines

Generation of a *chchd10* and a *chchd2* KO zebrafish models was performed using previously described (43). Briefly, Cas9 mRNA (100 ng/μl) and gRNA (100 ng/μl) were co-injected into embryos, using a volume of 1.5 nl, at the one cell stage of development. Founder zebrafish were identified at adult stages by screening their progeny for germline transmission of indels. From the offspring of candidate founders, DNA was extracted from 24 embryos, aged 24 hrs post fertilization (hpf), and polymerase chain reaction (PCR) amplification of the target site of interest was carried out followed by restriction fragment length polymorphism (PvuII and Pst1 for *chchd10* and *chchd2* respectively). Sanger sequencing (Genome Québec) was performed on amplicons to determine the exact mutation of each founder line. Primers sequences for verification of *chchd10* KO line: forward primer: GGGGGAAAAAAACAGTTCTTGGC; reverse primer: AAGAGTTGTTGGTTACACTCCAT. Primers sequences for verification of *chchd2* KO line: forward primer: AAAGGACAGGTAATTGATTGTGC; reverse primer: GCCATAACGTTTACCTGATAGGT.

### Larval Morphology, Motor Function and Survival Assays

To determine whether the mutants had any visible morphological defects, dechorionated 56 hpf larvae were examined. Heart defects, including asystole, bleeding, or edema (swelling in heart region) were quantified as previously described (44). Reduced yolk sack size was counted when the larvae’s yolk sack diameter was smaller than the diameter from embryo head (10 μm).

Touch-evoked motor response was evaluated between 54–56 hpf as previously described (45). Briefly, larvae were placed in the middle of a circular arena (150 mm diameter petri dish) containing fresh system water maintained at a temperature between 27-28.5 °C. A single touch to the tail was applied to larvae to initiate burst swimming behaviour and movements were recorded from above (recorded at 30 Hz; Grasshopper 2 camera, Point Grey Research). Swim distance, mean and maximum swim velocities were quantified using the manual tracking plugin for ImageJ.

Free swimming velocity (mm/s) was examined using motion tracking software (DanioVision, Noldus) in larvae aged 5 days post fertilization (dpf) placed in 24-well plates. Following a 5 min habituation period, motor activity was recorded for a total of 30 min. System water temperature was maintained at 28.5 °C. For survival assays, forty recently hatched 5 dpf larvae from each genotype were placed in a temperature-controlled container (28.5 °C) and monitored from 6 to 30 dpf. Water changes were made once every 2 days.

### RNA Extraction and Reverse Transcription–qPCR (RT-qPCR)

RNA was extracted from 5 pooled larvae aged 5 dpf using the PicoPure RNA Isolation Kit (Applied Biosystems). After quality control, 1 μg was used to create a cDNA library by reverse transcription using SuperScript VILO cDNA Synthesis Kit (Invitrogen). Quantitative Real time PCR was performed using SYBR Green Supermix (Bio-Rad laboratories). Relative quantification was performed using the delta-delta Ct method (46) with *actin* and *sdha* as internal controls (see supplementary material for primer sequences).

### Neuromuscular Junction Co-Localization

To investigate whether neuromuscular junction (NMJ) integrity was affected, double-labelling was performed using synaptotagmin 2 (Syt2, presynaptic marker) and alpha-bungarotoxin (*α*Btx, post-synaptic marker) as described before (45). Briefly, 2 dpf larvae were fixed with 4 % paraformaldehyde in Phosphate-buffered saline (PBS) overnight at 4°C with gentle rotation. Larvae were rinsed with PBS (4 times 15 min), followed by a 45-min incubation at room temperature (RT) with gentle rotation in PBS containing 1 mg/ml collagenase to remove skin. Larvae were then rinsed with PBS (4 times 10 min) and incubated for 30 min in PBST (PBS;Triton X-100) at RT with gentle rotation. Larvae were then incubated in 10 mg/ml sulforhodamine conjugated *α*Btx in PBST for 30 min at RT with gentle rotation. Larvae were then rinsed (4 times 10 min, PBST) and incubated in blocking solution (2% Goat Serum, 1% BSA, 0.1% TritonX-100, 1% DMSO in PBS) for 1 hour (hr) at RT. Primary antibody Syt2 (1:100, DSHB in blocking solution) was applied overnight at 4°C and gentle rotation. The samples were washed (4 times 30 min, PBST) and then incubated with the secondary antibody (Alexa fluor 647, 1: 1200 ThermoFisher in blocking solution) overnight at 4°C with gentle rotation. Larvae were rinsed (4 times 20 min PBST) and mounted on a glass slide in 70% glycerol. The NMJs were visualized with 60x/1.42 oil immersion objective. Images were acquired using the Volocity software (Improvision). Co-localization analysis was performed by determining the number of orphaned receptors, both pre- and post-synaptic, present in individual ventral roots.

### Immunoblot, Blue Native PAGE, and Second Dimension PAGE

For Immunoblots, larvae aged 5 dpf were deyolked using modified Ca^2+^ free Ginsburg Fish Ringer (55 mM NaCl, 1.8 mM KCl, 1.25 mM NaHCO_3_) solution. For each condition, 10 larvae were homogenized at 4 °C using RIPA buffer (50 mM Tris-HCl, pH 7.4, 100 mM NaCl, 1% NP40, 0.1 % SDS, 0.5 % sodium deoxycholate) and a protease inhibitor cocktail (cOmplete ULTRA Tablets Mini EDTA-free Protease Inhibitor Cocktail, Roche). Lysates were recovered following centrifugation (20 min at 12,000 rpm). Protein concentrations were quantified using a Bradford assay and 30 μg of lysate was separated on a 12.5% SDS polyacrylamide gel or a 4-16% gradient gel. Proteins were transferred to a nitrocellulose membrane and blocked in 5% milk (TSBT) for 1 hr at RT. Membranes were then incubated with primary antibodies in blocking buffer at 4 °C overnight followed by respective secondary antibody labelling (Jackson Immuno Research Laboratories, 1:10 000) for 1 hr at RT. Enhanced chemiluminescence (Clarity™ Western ECL Substrate, Bio-Rad) was used to visualize immunolabelled proteins. The following antibodies were used: Nudfa9 (ab14713, Abcam), Sdhb (ab14714, Abcam), Uqcrc2 (ab14711, Abcam) Mt-co1 (ab14705, Abcam), Atp5a (ab14748, Abcam), Vinculin (V4505, Sigma), Actin (LLC 691001, MP Biomedicals), Chchd10 (ab121196, Abcam), and a custom Chchd2 antibody (MédiMabs) targeting the following epitope sequence: PDVTYQEPYQGQAM. For blue native PAGE, one 5 dpf larvae was placed in MB2 buffer (0.5 ml of 3XGB (1.5 m aminocaproic acid, 150 mM Bis-Tris, pH: 7.0), 0.5 ml of 2M aminocaproic acid, 4 µl of 500 mM EDT) and homogenized at 4 °C. Lauryl maltoside was then added to a final concentration of 1% and incubated for 15 min at 4 °C. Following centrifugation, the supernatant was recovered and ran on a non-denaturing polyacrylamide 6-13% gradient gel. Separated complexes were transferred to a PVDF membrane. Membranes were blocked and probed as described above. Sdhb or Complex II (blue native PAGE) were used as a mitochondrial loading controls and actin was used as a cytoplasmic loading control. For the two-dimensions PAGE, strips were cut off from a first-dimension native gel that contained 30 μg of protein from 5 larvae of respective genotypes. Strips were then incubated for 45 min in 1 % SDS and 1% β-mercaptoethanol and run individually on a 10% SDS-PAGE to separate proteins, as described before. Quantifications of protein bands were performed using ImageJ.

### Pharmacology

Dechorionated 24 hpf larvae were treated with 100 mM 2-deoxy-d-glucose (2-DG, sigma D8375-5G diluted in E3 buffer (5 mM NaCl 0.17 mM KCl 0.33 mM CaCl_2_ 0.33 mM MgSO4, pH. 7.2) to inhibit glucose metabolism for 24 hr. Five treated and untreated larvae were then pooled and submitted to quantitative PCR or Immunoblot analysis as described above.

### Statistical Analysis

All statistical analyses were performed using Prism 7 (GraphPad Software Inc.). Two samples were compared using unpaired students’ t tests. Shapiro-Wilks test was used to test for normality. Kruskal-Wallis tests were used to compare more than 2 samples that were not normally distributed, followed by Dunn’s post hoc multiple comparisons test. One-way ANOVA and Tuckey’s multiple comparison were used when data was normally distributed. Survival curves were compared using the log rank test, and chi-square tests were used to compare proportions. Significance was assessed at *p* < 0.05.

## Results

### Generation of *chchd10 ^-/-^*, *chchd2 ^-/-^*, and double *chchd10 ^-/-^* & *chchd2 ^-/-^* zebrafish lines

A previous study investigating the role of Chchd10 in zebrafish using a morpholino to knockdown expression of the *chchd10* transcript showed that loss of *chchd10* resulted in reduced motor activity and survival, as well as altered myofibrillar structure and ventral root length (33). To avoid the variability and off-target effects associated with morpholino injections we generated genetic KO models of both *chchd10* and *chchd2* using CRISPR/Cas9 editing. We hypothesized that KO models would permit investigations further into development, as dependence on mitochondrial oxygen consumption for survival has been documented to increase with development in zebrafish (47, 48).

In zebrafish, Chchd10 and Chchd2 are encoded by genes containing 4 exons (Fig. 1 A, B). We selected guide RNAs that targeted the coding region of exon 2 in each gene and identified a single nucleotide insertion in *chchd10* and a seven base pair deletion in *chchd2* (Fig. 1 C, D). Both indels were transmitted to the next generation of offspring and could be identified by loss of PvuII and PstI restrictions sites respectively (Fig. 1 E, F, G, H). To investigate whether our mutations disrupted gene function in each fish line, we quantified expression of each transcript by RT-qPCR from 5 dpf larvae and observed a significant reduction in *chchd10* and *chchd2* expression levels in homozygous mutant carriers, suggesting that the transcripts were subjected to nonsense-mediated decay (Fig 1 Ii, Ji). Immunoblot analysis confirmed the complete loss of Chchd10 and Chchd2 protein expression in homozygous carriers (Fig. 1 Iii, Jii). A double *chchd10*^-/-^ & *chchd2*^-/-^ model was then generated by in-crossing the two single mutant lines.

**Figure 1.**
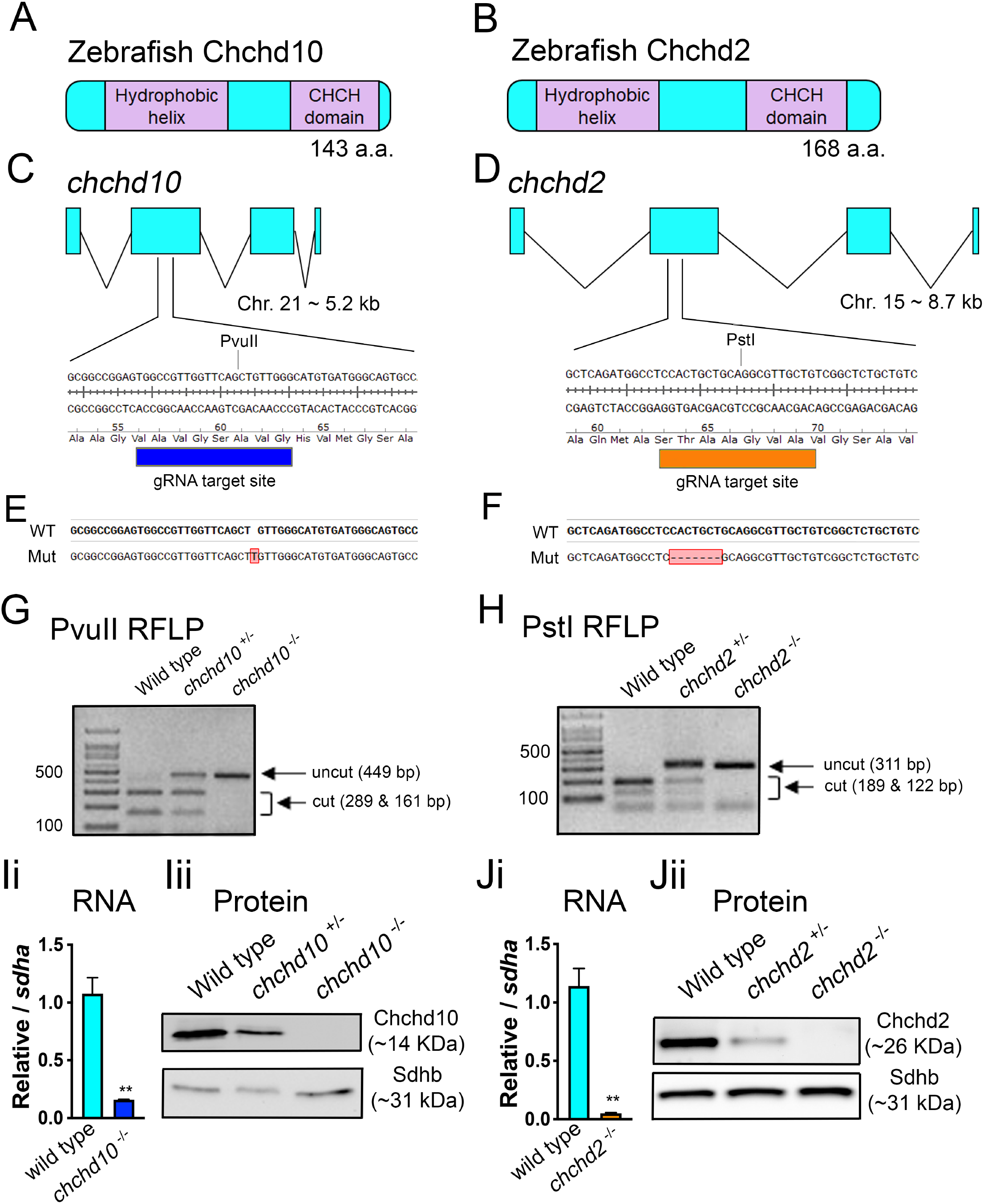
Generation of *chchd10 ^-/-^* and *chchd2 ^-/^*^-^ zebrafish lines. Schematic representation of the zebrafish proteins Chchd10 (**A**) and Chchd2 (**B**). **C**, *chchd10* gene structure and gRNA target site. **D**, *chchd2* gene structure and gRNA target site. **E**, CRISPR-induced *chchd10 ^-/-^* mutant (mut) line. **F**, CRISPR-induced *chchd2 ^-/-^* mutant (mut) line. **G**, Restriction fragment length polymorphism (RFLP) screening method using PvuII which is lost in the *chchd10 ^-/-^* mutant. **H**, RFLP screening using PstI which is lost in the *chchd2 ^-/-^* mutant. **Ii**, RT-qPCR of the *chchd10* transcript using wild type and *chchd10 ^-/-^* cDNA. A reduction in *chchd10* (t (4) = 6.61, *p* = 0.003, relative to *sdha*) was observed in *chchd10 ^-/-^* (n= 3 per genotype, data represent mean ± S.E.M). **Iii**, Immunoblot using wild type, *chchd10 ^+/-^*, and *chchd10 ^-/-^* whole brain lysate. Sdhb was used as a loading control. **Ji**, RT-qPCR of the *chchd2* transcript from wild type and *chchd2 ^-/-^* cDNA. A significant reduction in *chchd2* (t (4) = 7.26, *p* = 0.002, relative to *sdha*) was observed in *chchd2 ^-/-^* (n= 3 per genotype, data represent mean ± S.E.M). **Jii**, Brain lysate immunoblot from wild type, *chchd2 ^+/ -^, chchd2 ^-/-^* model. Sdhb was used as a loading control.

### Larval *chchd10^-/-^*, *chchd2^-/-^*, and double *chchd10^-/-^*& *chchd2^-/-^* display morphological defects, motor impairment, reduced survival, and altered neuromuscular integrity

Morphological observations of larvae at 56 hpf showed ∼64% of *chchd10*^-/-^ larvae had a reduced yolk sac size compared to wild type larvae, while *chchd2*^-/-^ and double *chchd10*^-/-^ & *chchd2*^-/-^ did not (Fig. 2 A, B). Cardiac defects were present in 37% of *chchd2*^-/-^ larvae and 30% of double *chchd10*^-/-^ & *chchd2*^-/-^ larvae, with most larvae displaying a cardiac edema (Supp. video 1); however, this phenotype was not observed in *chchd10*^-/-^ larvae (Fig. 2 A, C).

**Figure 2.**
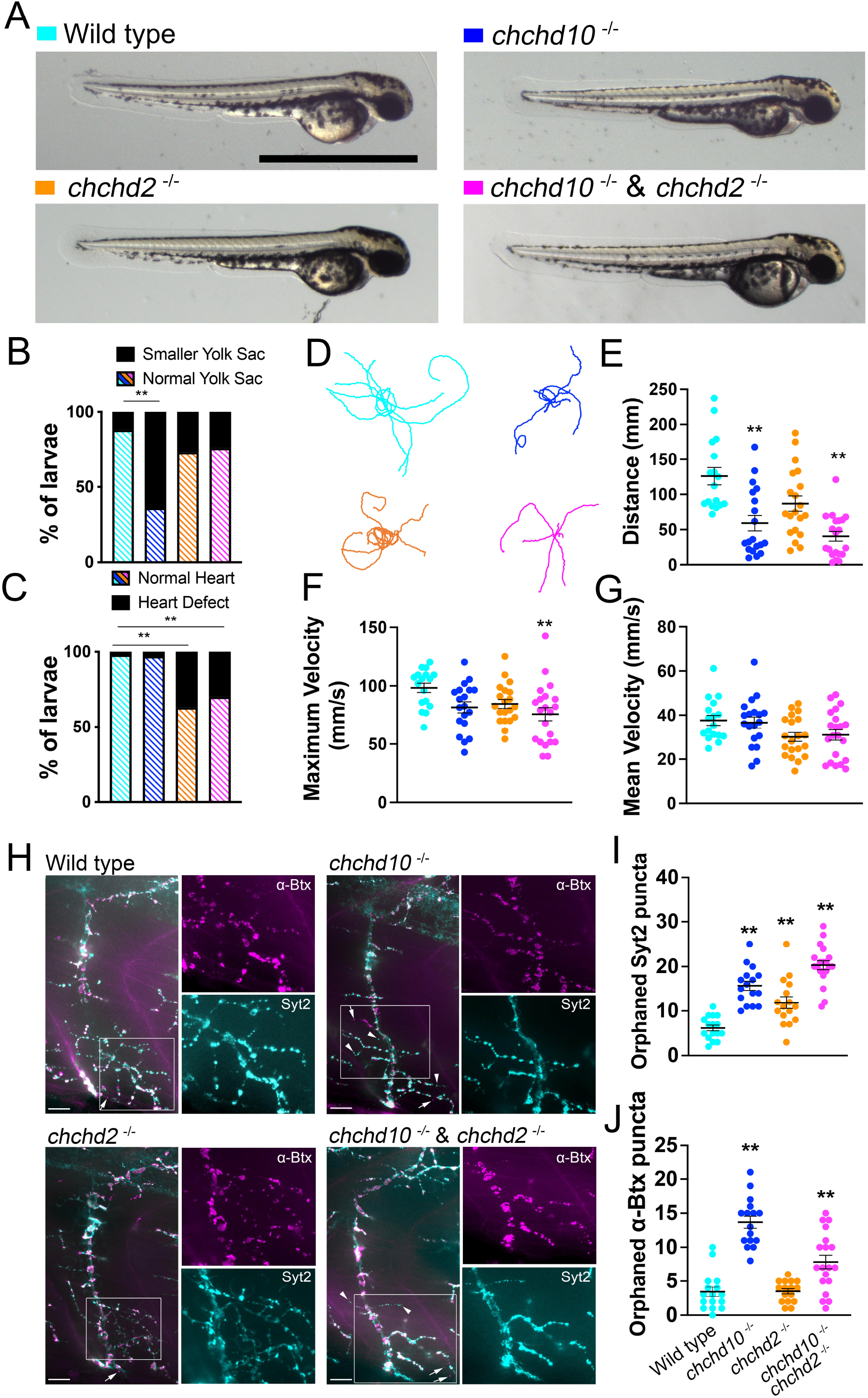
Larval *chchd10*^-/-^, *chchd2*^-/-^ and double *chchd10*^-/-^ & *chchd2*^-/-^ display morphological defects, motor impairment and altered neuromuscular junction integrity. **A,** Representative images of 56 hpf larvae. **B**, Quantification of yolk sacs defects. Larval *chchd10*^-/-^ showing reduced yolk sack size compared to wild type larvae (*χ*^2^= 18.5, *p* < 0.0001). Sample sizes were as follows: wild type, n = 34; *chchd10*^-/-^, n = 28; *chchd2*^-/-^, n = 30, and double *chchd10*^-/-^ & *chchd2*^-/-^, n = 37. **C**, Quantification of heart defects. Both *chchd2*^-/-^ (*χ*^2^= 23.43, *p* < 0.0001) and double *chchd10*^-/-^ & *chchd2*^-/-^ (*χ*^2^= 16.05, *p* < 0.0001) larvae display cardiac abnormalities. Sample sizes are as follows: wild type, n = 58; *chchd10*^-/-^, n = 35; *chchd2*^-/-^, n = 62, and double *chchd10*^-/-^ & *chchd2*^-/-^, n = 37. **D**, Representative traces of touch-evoked motor response 2 dpf larvae (n =10 per genotype). Quantification of swim distance (**E**), maximum swim velocity (**F**), and mean swim velocity (**G**). Data shown as individual data points ± S.E.M. Samples sizes were as follows: wild type, n = 17; *chchd10*^-/-^, n = 19; *chchd2*^-/-^, n = 20, and double *chchd10*^-/-^ & *chchd2*^-/-^, n = 20. Significance was assed using Kruskal-Wallis followed by Dunn’s post hoc test for swimming distance and by one-way ANOVA and Tuckey’s multiple comparison test for maximum velocity and mean distance measures. **H**, Representative images of NMJs in the ventral trunk musculature of larvae aged 2 dpf. Markers are Syt2 (cyan) and *α*Btx (magenta). Arrow and arrowheads in example NMJ images represent orphaned *α*Btx and Syt2 puncta, respectively. Scale bars represent 100 µm. **I**, Tabulation of orphaned pre-synaptic (Syt2) puncta. All genetic groups were significantly different than wild type NMJs (F (3, 63) = 34.33, *p* < 0.01). Data shown as mean ± S.E.M. **J**, Tabulation of orphaned post-synaptic (*α*Btx) puncta. Both *chchd10 ^-/-^* and double *chchd10 ^-/-^ & chchd2 ^-/-^* showed increased orphaned AChR clusters when compared to wild type larvae (F (3, 63) = 34.80, *p* < 0.01). Data shown as mean ± S.E.M Sample sizes are as follows for both presynaptic and post-synaptic quantification: wild type, n=16 ventral roots, 6 larvae; *chchd10 ^-/-^*, n=16 ventral roots, 6 larvae; *chchd2 ^-/-^*, n=16 ventral roots, 6 larvae, and double *chchd10 ^-/-^* & *chchd2 ^-/-^*, n=19 ventral roots, 7 larvae. Significant difference with wild type larvae is represented by a single asterisk (*p* < 0.05) or double asterisk (*p* < 0.01).

As ALS impairs motor function, we next examined locomotor function in our zebrafish models by examaning touch response behaviour at 2 dpf. Touch-evoked motor responses were recorded and changes in swimming behaviour were observed in some of our measures of motor function (see 10 representative traces per genotype, Fig. 2 D). Specifically, mean swimming distance was significantly reduced in *chchd10*^-/-^, and double *chchd10*^-/-^ & *chchd2*^-/-^ larvae when compared to wild type zebrafish. The *chchd10*^-/-^ larvae swam half of the distance of wild type larvae, while the double *chchd10*^-/-^ & *chchd2*^-/-^ larvae only swam a third of the distance swam by wild type larvae (Fig. 2 E). A significant difference was also observed in the maximum velocity measure; double *chchd10*^-/-^ & *chchd2*^-/-^ larvae swam at a lower maximum velocity compared to wild type (Fig. 2 F), while no significant differences were observed in mean velocity (Fig. 2 G).

To determine whether the early motor defect could be due to reduced neuromuscular junction (NMJ) integrity as has been described in other ALS models (45, 49, 50), we explored co-localization of pre-synaptic and post-synaptic markers at 2 dpf. While few orphaned post- and pre-synaptic markers were observed in wild type larvae, the mutant models showed significant orphaned clusters of both pre-synaptic (syt2) and post-synaptic markers (*α*Btx). All mutants showed increased orphaned pre-synaptic markers compared to wild type (Fig. 2 H, I), while only *chchd10*^-/-^ and double *chchd10*^-/-^ & *chchd2*^-/-^ had increased orphaned post-synaptic markers compared to wild type larvae (Fig. 2 H, J).

Measures of locomotor function were then performed on 5 dpf larvae to see whether defects worsened during development due to increased reliance on oxygen consumption. Larvae were individually placed in 24-well plates filled with fish-system water and traces were recorded to measure mean swim velocity over a 30-min period from animals in all four genetic groupings (Fig. 3 A). At that age, mean swim velocity was significantly reduced in all our models, with the double *chchd10*^-/-^ & *chchd2*^-/-^ showing the greatest deficit (Fig. 3 A, B).

**Figure 3.**
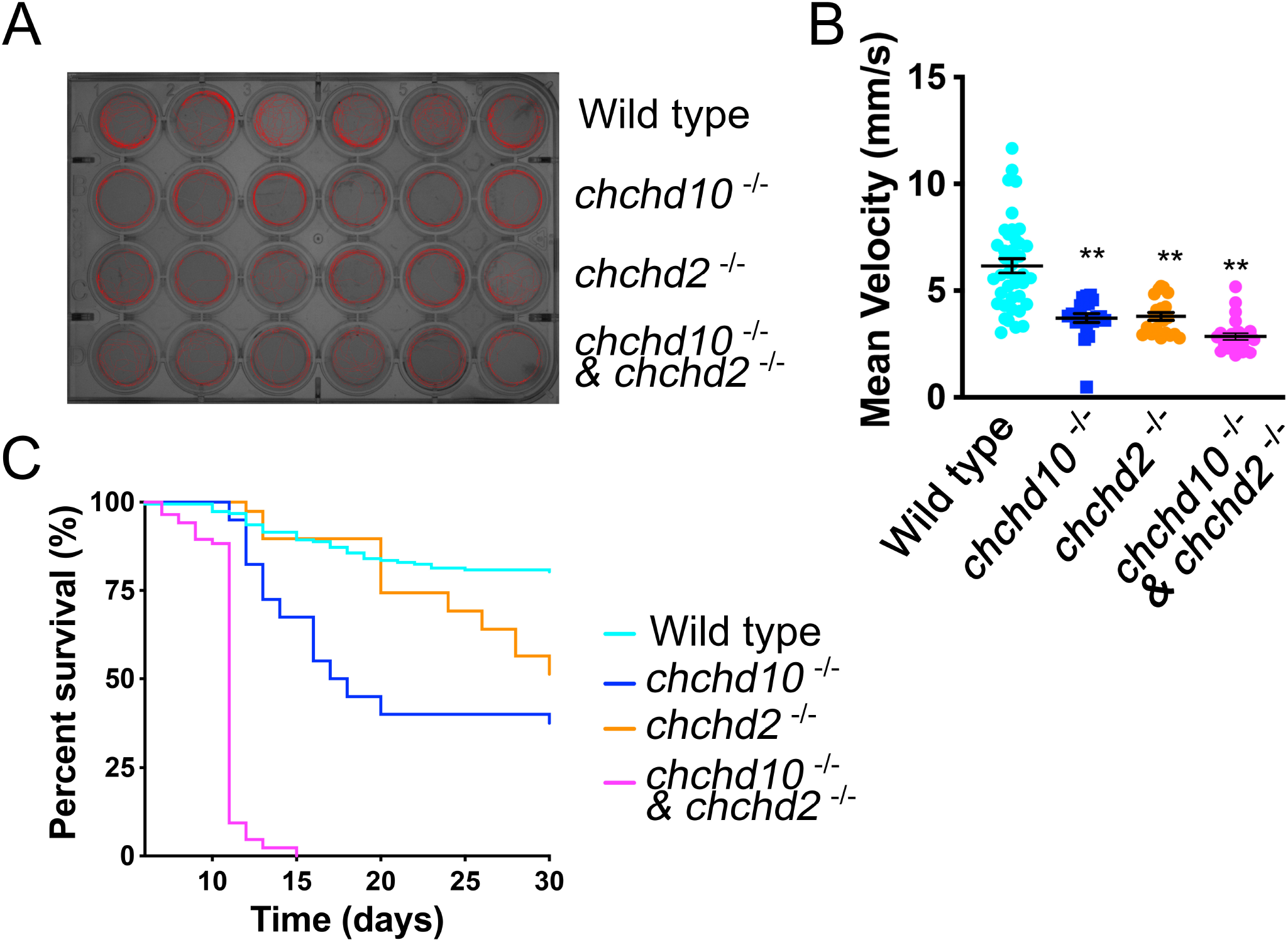
All mutant larvae present a motor deficit at 5 dpf and reduced survival. **A**, Representative image of a 24-well plate free swim arena and movement paths (6 per genonytpe) of larvae aged 5 dpf. **B**, Quantification of mean velocity. Specific mean swim velocities are as follows: wild type, 6.2 mm/s, n = 40; *chchd10^-/-^*, 3.72 mm/s, n = 21; *chchd2^-/-^*, 3.80 mm/s, n = 21, and double *chchd10*^-/-^ & *chchd2*^-/-^, 2.87 mm/s, n = 36. Data represented as individual data points ± S.E.M. Kruskal-Wallis test was used to assess significance. Double asterisks represent *p* < 0.01, Dunn’s Post hoc test. **C**, Survival rates were significantly reduced in all mutants, with no double *chchd10* ^-/-^ & *chchd2* ^-/-^ surviving past 15 days. Sample sizes are as follows: wild type, n =189; *chchd10*^-/-^, n = 40; *chchd2*^-/-^, n = 39, and double *chchd10*^-/-^ & *chchd2*^-/-^, n = 86. Significance was assessed using Mantel-Cox test, all *p* <0.01 compared to wild type.

We next examined larval survival and observed reduced survival for *chchd10*^-/-^ (median survival 18 days) and *chchd2*^-/-^ (did not reach median survival) models when compared to wild type larvae. Under these rearing conditions, no double *chchd10*^-/-^ & *chchd2*^-/-^ fish survived past 15 dpf (median survival 11 days) (Fig. 3 C).

### Chchd10 compensates for loss of Chchd2 but not *vice versa,* and forms both a low and high molecular weight complex at 5 dpf

We next explored whether the KOs of Chchd10 and Chchd2 elicited compensatory changes in expression of either protein or the formation of higher molecular weight complexes (9, 10). Using lysates from 5 dpf larvae, we performed Immunoblots and observed an increase (1.9 fold) in Chchd10 protein levels in our *chchd2*^-/-^ zebrafish line compared to wild-type (Fig. 4 Ai, Aii). Unexpectedly, Chchd2 protein levels in *chchd10*^-/-^ lysates were 80 % lower compared to wild type lysate (Fig. 4 Bi, Bii). We next examined whether there were changes in their transcript levels in the KO models. No significant differences in the *chchd10* transcript levels in the *chchd2*^-/-^ model, nor any change in the *chchd2* transcript levels in the *chchd10* ^-/-^ model were observed when compared to wild type larvae (Fig. 4 Aiii, Biii), suggesting that the changes in protein levels likely occur post-transcriptionally.

**Figure 4.**
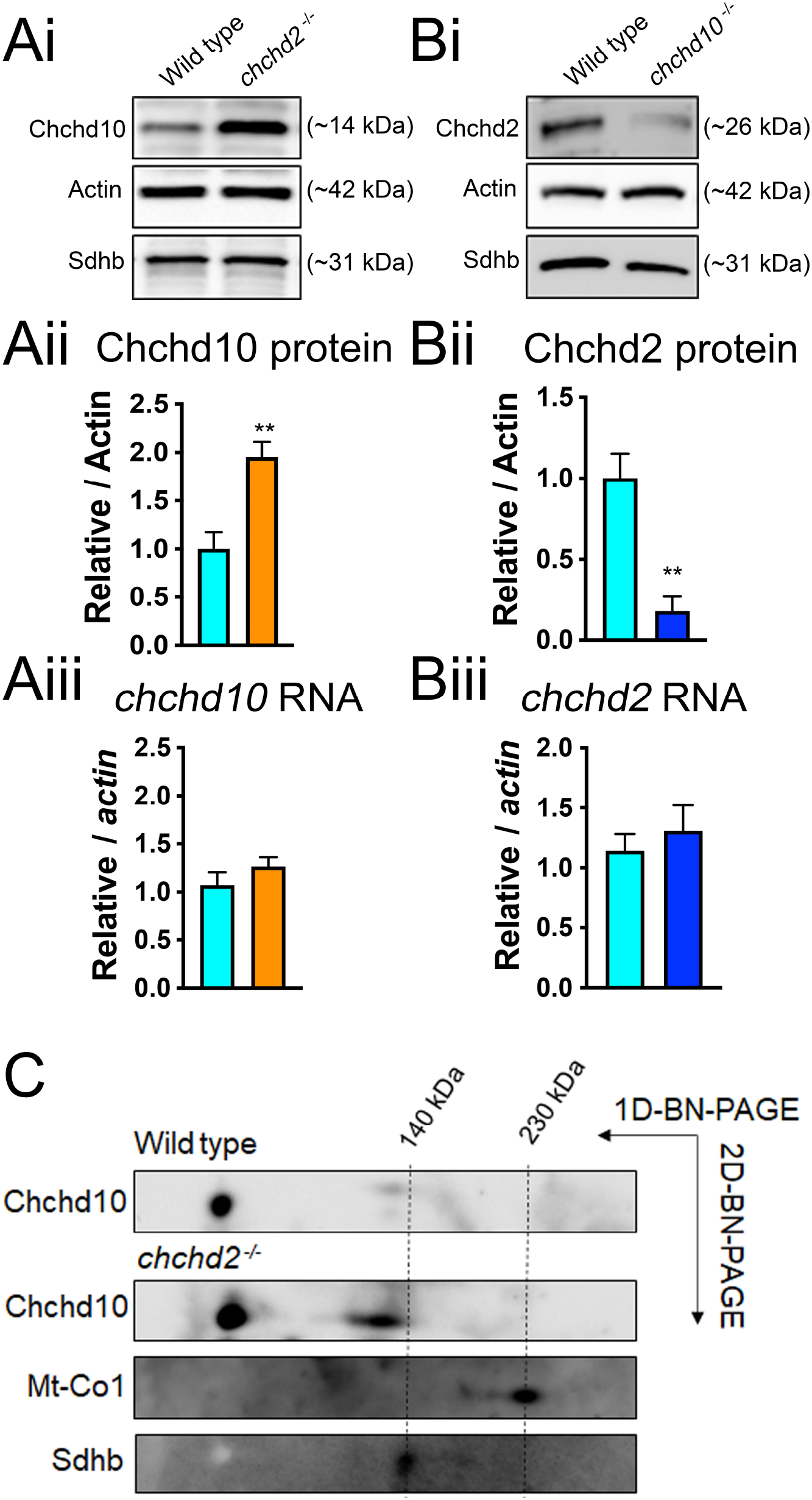
Larval Chchd10 compensates for loss of Chchd2 expression but not *vice versa* and is present in both a lower and higher molecular weight complex in zebrafish at 5 dpf. **Ai**, Representative immunoblot of Chchd10 protein level in *chchd2*^-/-^ larvae aged 5 dpf. **Aii**, Immunoblot quantification (n=3 per genotype, 5 dpf) showing increased Chchd10 expression compared to wild type in *chchd2*^-/-^ larvae (*t* (4) = 4.04, *p=*0.011). Actin was used as loading control. **Aiii**, Unchanged transcript levels of *chchd10 in* 5 dpf *chchd2*^-/-^ larvae (*t* (4) = 0.30, *p* = 0.30 relative to *actin*). **Bi**, Representative immunoblot of Chchd2 protein expression in *chchd10*^-/-^ larvae aged 5 dpf. **Bii,** Immunoblot quantification (n=3 per genotype, 5 dpf) showing reduced Chchd2 expression levels in *chchd10*^-/-^ larvae compared to wild type larvae (*t* (4) = 4.62, *p=*0.0099). Actin was used as loading control. **Biii**, Unchanged transcript levels of *chchd2* in 5dpf *chchd10*^-/-^ larvae (*t* (4) = 0.55, *p* = 0.55, relative to *actin*). **C**, Second dimensional PAGE analysis using a Chchd10 antibody revealed a high molecular weight complex, close to 140 kDa, in both wild type and *chchd2*^-/-^ larvae. Mt-Co1 and Sdhb are used as molecular weight markers.

To investigate whether Chchd10 and Chchd2 formed a stable high molecular weight complex as previously described in cellular models (10), a two-dimensional gel (2D-gel) analysis was performed on lysates collected from our genetic groupings aged 5 dpf. In this analysis, protein complexes are first separated on a blue native gel, and then a gel strip containing the separated complexes is run on a denaturing gel to detect proteins within complexes of various molecular weights. While our Chchd2 antibody was not able to detect the protein on the 2D-gel, our Chchd10 antibody revealed that Chchd10 is present mostly as a low molecular weight species (monomer or dimer), as in cellular models (10) (Fig. 4 C). The presence of a higher molecular weight complex in wild type and *chchd2*^-/-^ lysate was also observed consistent with observations in cellular models (Fig. 4 C), though the complex was ∼140 kDa in our model *vs* ∼170 and 230 kDa in human fibroblasts (10). Finally, elevated levels of both the low and higher molecular weight Chchd10 complex were observed in the *chchd2*^-/-^ lysate (Fig. 4 C).

### Chchd10 and Chchd2 protein levels increase in response to forced reliance on mitochondrial energy production in wild type larvae

We next examined whether Chchd10 and Chchd2 levels were responsive to mitochondrial stressors as described previously (28). Wild type larvae aged 24 hpf were exposed to the glycolytic inhibitor 2-DG for 24 hr, permitting us to test whether Chchd10 and Chchd2 expression is affected when fish are forced to rely on mitochondria for energy production (Fig. 5 A). Immunoblot analysis revealed that exposure to 2-DG led to increased expression of Chchd10 and Chchd2 (Fig. 5 B, Ci, Cii). Exposure to 2-DG did not result in a significant change of *chchd10* or *chchd2* transcript levels showing that the increase in both proteins was regulated post-transcriptionally (Fig. 5 Cii, Civ). Interestingly, touch-evoked motor behaviour analysis in 2 dpf larvae revealed a concomitant motor deficit on exposure to 2-DG. More specifically, we observed reduced swim distance in wild type larvae exposed to 2-DG while maximum swim velocity was unchanged when compared to wild type larvae not treated with 2-DG (Fig. 5 Di, Dii, Diii).

**Figure 5.**
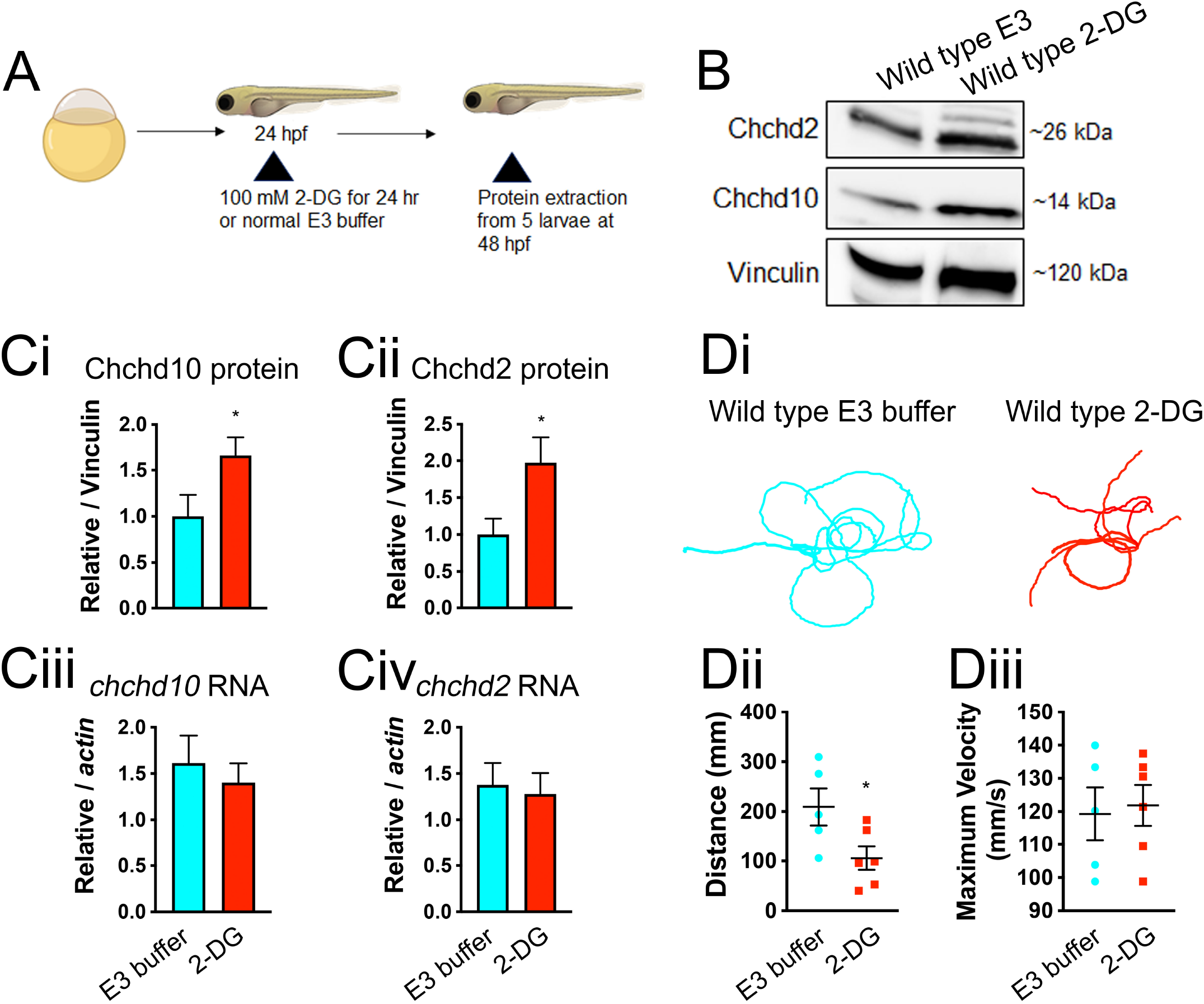
Both Chchd10 and Chchd2 proteins display elevated expression levels in response increased mitochondrial reliance in wild type larvae. **A**, Experimental timeline for 2-DG treatment (Images were created using BioRender). **B**, Example immunoblot showing Chchd10 and Chchd2 expression change following treatment with to 2-DG. **Ci, Cii**, Quantification of immunoblot data. Significant difference in both Chchd10 (*t* (10) = 2.184, *p* = 0.053), and Chchd2 (*t* (10) = 2.348, *p* = 0.04) protein levels following 2-DG treatment was observed. Sample sizes are as follows: 2-DG, n = 6; E3 Buffer, n = 6. Vinculin was used as loading control. **Ciii, Civ**, Unchanged expression levels following 2-DG exposure in both transcripts: *chchd10* (*t* (4) = 0.59, *p* = 0.59, relative to *actin*) and *chchd2*: (*t* (4) = 0.78, *p* = 0.78 relative to *actin*). Sample sizes are as follows: 2-DG, n = 3; E3 buffer, n = 3. Single asterisks represent significant differences from E3-treated wild type (*p* < 0.05). **Di**, Representative touch-evoked larval motor responses following exposure to 2-DG or E3 buffer treatment for 24 hrs. Motor responses were evaluated at 56 hpf. **Dii**, Larvae exposed to 2-DG show significantly reduced swim distance (*t* (9) = 2.45, *p* = 0.04), though maximum swim velocity. **Diii**, was not found to be significantly different (*t* (9) = 0.26, *p* = 0.80). Sample sizes as follows E3 Buffer, n =5; 2-DG, n = 6.

### Reduced assembly of mitochondrial Complex I in *chchd10^-/-^* and *chchd2^-/-^* zebrafish larvae, but not in double *chchd10^-/-^* & *chchd2^-/-^* larvae at 5 dpf

As all our knockout lines displayed a significant defect in motor function at 5 dpf compared to 2 dpf, we next investigated whether mitochondrial respiratory chain integrity was compromised at that stage of development. Immunoblots revealed a significant reduction of the Ndufa9 subunit (Complex I) in *chchd10*^-/-^ whole lysate but not in *chchd2*^-/-^ or double *chchd10*^-/-^ & *chchd2*^-/-^ lysate at 5 dpf. Subunits of other respiratory chain complexes did not show differences between wild type, *chchd10*^-/-^, *chchd2*^-/-^ or double *chchd10*^-/-^ & *chchd2*^-/-^ (Fig. 6 A, B). As Ndufa9 was reduced in the *chchd10*^-/-^ model, we explored whether Complex I assembly was affected by blue native PAGE analysis of single zebrafish larvae aged 5 dpf (Fig. 6 C). We observed a significant reduction of Complex I assembly, particularly in *chchd10*^-/-^, as well as in our *chchd2*^-/-^ larvae at 5 dpf. Surprisingly, this defect was not observed our double *chchd10*^-/-^ & *chchd2*^-/-^ model (Fig. 6 D), suggesting that the Complex I assembly deficit is not solely responsible for the observed changes in motor behaviour at 5 dpf.

**Figure 6.**
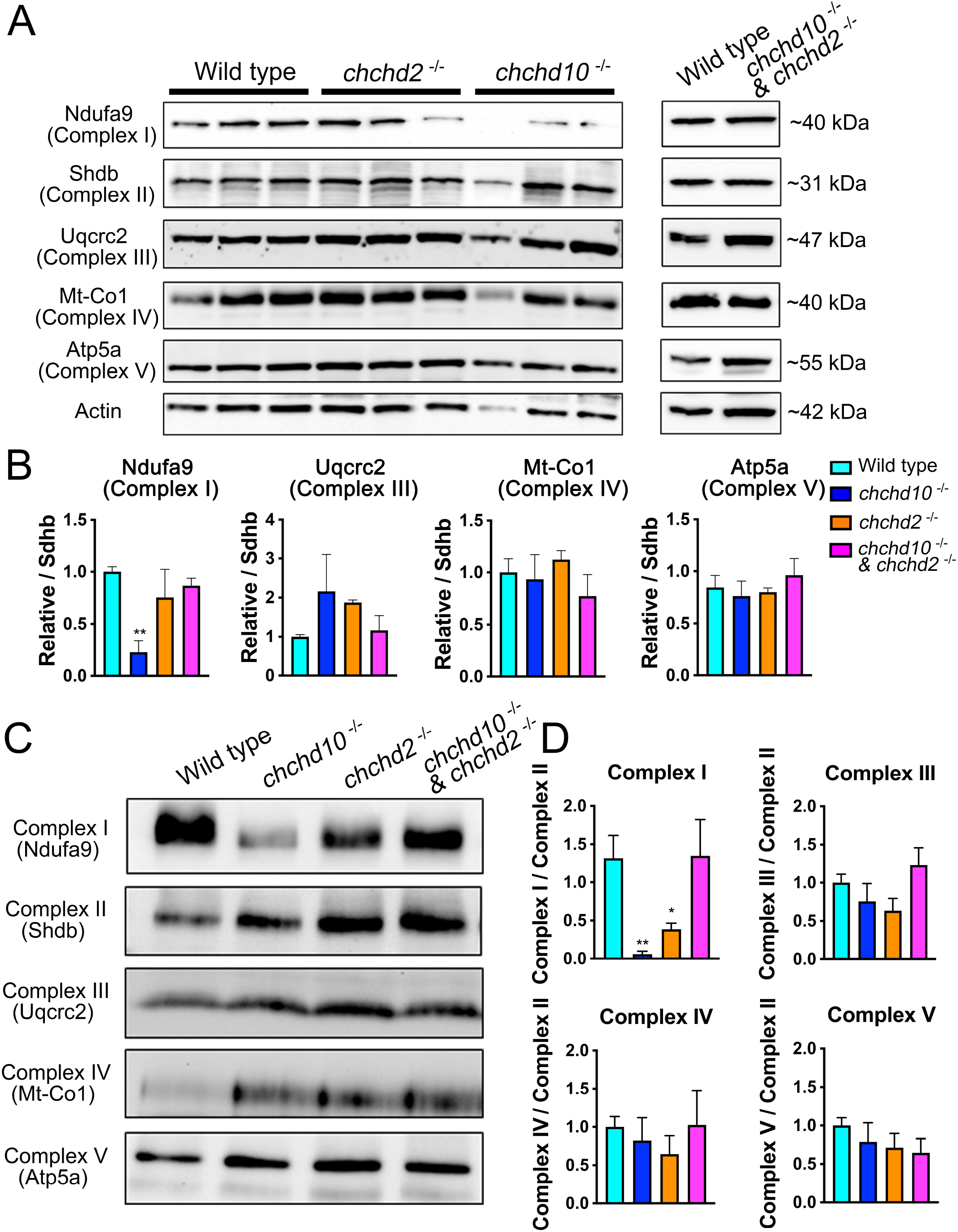
Reduced assembly of mitochondrial Complex I in *chchd10*^-/-^ and *chchd2*^-/-^ zebrafish larvae, but not in double *chchd10*^-/-^ & *chchd2*^-/-^ larvae at 5 dpf. **A**, Immunoblot analysis of mitochondrial respiratory chain subunits in larvae aged 5 dpf and quantification (**B**). Ndufa9 is significantly reduced in *chchd10 ^-/-^* larvae (F (3, 11) = 7.40, p = 0.006, *p* = 0.003 Tuckey’s, n = 3 per genotype). Data is represented as mean ± S.E.M. **C**, Representative blue native PAGE with each lane constituted of an individual 5 dpf larvae of respective genotype and quantification (**D**). Significant reduction of Complex I assembly was observed (F (3, 25) = 7.42, *p* < 0.01) in *chchd10*^-/-^ larvae (*p* < 0.01, Tuckey’s), and *chchd2*^-/-^ larvae (*p* < 0.05 Tuckey’s), but not in double *chchd10*^-/-^ & *chchd2*^-/-^ model. Sample size are as follows, wild type, n= 8; *chchd2*^-/-^, n = 7; *chchd10*^-/-^, n = 8, and *chchd10*^-/-^ & *chchd2*^-/-^ n=5. Data is represented as mean ± S.E.M. Significant difference with wild type larvae is represented by a single asterisk (*p* < 0.05) or double asterisk (*p* < 0.01).

### Markers of the mitochondrial integrated stress response are upregulated in *chchd10^-/-^* & *chchd2 ^-/-^* larvae at 5 dpf

Since no defect in mitochondrial respiratory chain complex assembly was observed in the double *chchd10*^-/-^ & *chchd2*^-/-^ model, we investigated whether other mechanisms could be underlying the observed deficits. As the mt-ISR has been documented in CHCHD10/2 models (18–20, 22), we explored whether the RNA levels of ISR markers were increased in the KO models at 5 dpf. While this pathway was not activated in the single gene KO models, a number of markers were increased in the double *chchd10*^-/-^ & *chchd2*^-/-^ model compared to wild type larvae. Transcripts orthologous to *ATF4* (*atf4a*, *atf4b*), *ATF5* (*atf5a*, *atf5b*) and CHOP (*chop*) were significantly increased in double *chchd10^-/-^* & *chchd2 ^-/-^*larvae, but not the single KOs (Fig. 7 A). *FGF21* and *HTRA2* orthologous genes (*fgf21*, *htra2*) were also significantly upregulated in the double *chchd10^-/-^* & *chchd2 ^-/-^* model (Fig. 7 B) as was the previously described oxygen responsive transcript, *cox4i2* (25–27) (Fig. 7 C).

**Figure 7.**
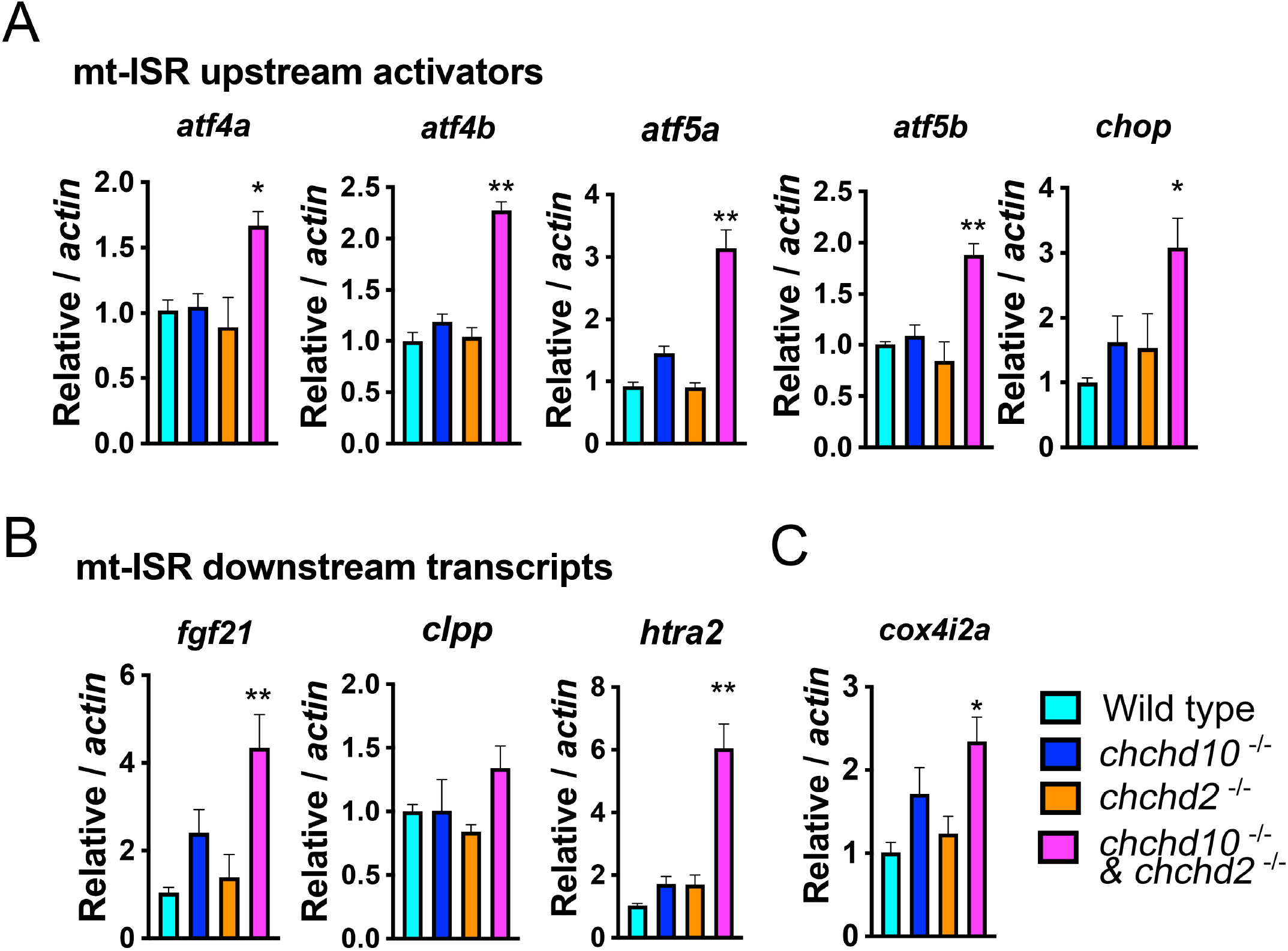
Several markers of the mitochondrial stress response are upregulated in our double *chchd10^-/-^* & *chchd2 ^-/-^* model at 5 dpf. **A**, All mt-ISR upstream activators were significantly increased in the double *chchd10^-/-^* & *chchd2 ^-/-^* model including *atf4a* (F (3, 8) = 5.964, *p* = 0.019); *atf4b* (F (3, 8) = 49.89, *p* <0.01); *atf5a* (F (3, 12) = 56.65, *p* < 0.01); *atf5b* (F (3, 8) = 14.39, *p* < 0.01); *chop* (F (3, 8) = 6.669, *p <* 0.01). **B**, Several downstream targets are upregulated in the double *chchd10^-/-^* & *chchd2 ^-/-^* model including *htra2* (F (3, 8) = 28.38, *p <* 0.01); *fgf21* (F (3, 8) = 9.177, *p<* 0.01), but not *clpp* (F (3, 9) = 2.032, *p* = 0.1799). **C**, Oxygen responsive transcript *cox4i2a* was significantly increased in our double *chchd10^-/-^* & *chchd2 ^-/-^* model (F (3, 7) = 6.919, *p* = 0.017). Data is represented as mean ± S.E.M, n = 3 per genotype. Significant difference with wild type larvae is represented by a single asterisk (*p* < 0.05) or double asterisk (*p* < 0.01), Tuckey’s multiple comparison test.

## Discussion

In this study, we demonstrate that loss of either or both Chchd10 and Chchd2 in zebrafish larvae leads to several ALS-like phenotypes including motor deficits, altered NMJ pre- and post-synaptic marker co-localization and reduced survival. As single gene KO mice generally express a very mild phenotype (28–30, 51), this suggests that zebrafish might be better suited for *in vivo* Chchd10 and Chchd2 loss of function studies. This study replicates *in vitro* findings in which CHCHD10 expression levels increase following loss of CHCHD2 (9, 10, 52); however, loss of Chchd10 led to reduced Chchd2 levels at 5 dpf. These data suggest that while the proteins are to some degree interdependent, they likely share only partially overlapping functions. Consistent with this, the double KO larvae displayed the most severe phenotypes.

A morphological heart defect, though not fully penetrant, was only observed in larvae that did not express Chchd2. Mitochondrial dysfunction is often linked to cardiomyopathies due to the dependence of cardiac function on OXPHOS (53), and heart defects result from exposure to various mitochondrial inhibitors in zebrafish (44). Interestingly, as double CHCHD10/CHCHD2 KO mice (21), but not CHCHD10 KO mice (19) die from a cardiac defect, it is possible that loss of Chchd2 has a greater role in conferring a cardiac defect than loss of Chchd10, as suggested by our study. Our study also highlighted that Chchd10 might be more involved in NMJ stabilization than Chchd2. While all mutants showed increased orphaned pre-synaptic NMJ markers, only *chchd10* ^-/-^ and double *chchd10* ^-/-^ & *chchd2* ^-/-^ displayed orphaned post-synaptic AchRs, which may account for the reduced motor function in our models at 2 dpf. A muscle conditional KO of CHCHD10 study in mice demonstrated that CHCHD10 was important for agrin-induced AchR clustering (54) which could lead to reduced organization and subsequently decreased post-synaptic innervation by ventral root projections as was observed in our CHCHD10 KO model.

To investigate potential mechanisms that might underlie the observed ALS-phenotypes, we examined the integrity of the OXPHOS system in our mutant models. Motor deficits at 5 dpf were accompanied by reduced Complex I assembly in both single gene KO models. This replicates *in vitro* results where reduced Complex I assembly was described in patient fibroblasts expressing the CHCHD10 R15L variant (10). Interestingly, the Complex I assembly defect in *chchd2*^-/-^ larvae was not accompanied by reduced Ndufa9 subunit expression. This suggests that compensation by Chchd10 can mitigate the subunit expression, but not the assembly defect, and highlights that Chchd2 may be more involved in OXPHOS complex assembly and/or stability, as has been previously proposed for Complex IV and to a lesser extent Complex I (55).

The most surprising finding of this study was that although the double *chchd10* ^-/-^ & *chchd2* ^-/-^ larvae displayed the most robust ALS-like phenotype, we did not observe a Complex I assembly defect. This suggests that reduced assembly of Complex I may not be solely responsible for the ALS-like phenotypes, but that other, more deleterious, cellular pathways are altered. The mt-ISR, a conserved transcriptional response that can be a triggered by mitochondrial dysfunction (56), has been linked to neurodegeneration (57) and has been described in CHCHD10/2 models previously (18–22). Transcripts for all three upstream mt-ISR activators (*atf4, atf5, chop*) (56), and *fgf21*, a biomarker of mitochondrial stress (58, 59) were significantly increased in our double *chchd10*^-/-^ & *chchd2*^-/-^ model, but not in the single KO models. This is in line with previous studies where activation of the mt-ISR was observed in both CHCHD10 S55L knock-in mice (19), double KO mice and cells (20, 21), but not in CHCHD10 KO mice. A recent study using CHCHD10 S55L mice suggested that mt-ISR precedes OXPHOS deficiency due to CHCHD10 protein aggregation (22). In contrast, our single gene KO models results suggest that OXPHOS deficiency is independent of the mt-ISR. Moreover, whether the OXPHOS deficiency triggers the mt-ISR, and whether the compensation of Complex I assembly in the double *chchd10*^-/-^ & *chchd2*^-/-^ is a downstream consequence of the activation of the mt-ISR, as has been suggested before in *C. elegans* (60), could be explored in future studies. Nevertheless, as the double *chchd10*^-/-^ & *chchd2*^-/-^ seem to have a worse phenotype than single gene KOs, this suggests that downstream consequences of the mt-ISR might be more detrimental than the observed Complex I deficiency in our single KO.

## Conclusion

In this study, we used three knockout models and showed that loss of either protein leads to distinct, but overlapping, phenotypes during zebrafish development. This supports the hypothesis that loss-of-function of either or both proteins might be involved in the pathogenicity of various neurodegenerative diseases and highlights that the biological mechanisms that confer disease still need to be explored. Most importantly, the recovery of Complex I defect in the double *chchd10*^-/-^ & *chchd2*^-/-^ as well as the activation of the mt-ISR suggests that this pathway, likely activated to resolve mitochondrial dysfunction, might lead to more detrimental consequences when activated in vertebrates.

## Supporting information

Heart Defect Video

Primer Sequences

Wild Type Heart Video

