## Supplementary material for "Loss of mitochondrial Chchd10 or Chchd2 in zebrafish leads to an ALS-like phenotype and Complex I deficiency independent of the mt-ISR": Primer Sequences

**Supplemental data:**

**Quantitative PCR primers:**

| | |
|---|---|
| *actin* forward | TGTTTTCCCCTCCATTGTTGG |
| *actin* reverse | TTCTCCTTGATGTCACGGAC |
| *sdha* forward | GAGTCTCCAATCAGTATCCAGTAGTAGA |
| *sdha* reverse | CACTGTGTGCGAGCGTGTTG |
| *chchd10* forward | CAGTTCAGAGGCACCCAAAC |
| chchd10 reverse | CACTGAGAGGGTGGAAGTCG |
| *chchd2* forward | GACATTCTGAAGCTGCTAGGC |
| *chchd2* reverse | CTGCTGTGGCGGGTACAT |
| *atf4a* forward | TGCACTGGCTCAGTTTGG |
| *atf4a* reverse | CGGAGAGATGGAGATGCTGT |
| *atf4b* forward | GGCACAGACAGTGGTGAAGA |
| atf4b reverse | AGAACTGGGTGGATCCTCAG |
| *atf5a* forward | CTCACCCACAGGCTAACCAC |
| *atf5a* reverse | CCAGTCACTAAGACCATCACCA |
| *atf5b* forward | AGCTTCTCCATTGCGGTTTA |
| *atf5b* reverse | CAGGAAAGAGATGAGAAAAACTGA |
| *chop* forward | CACAGACCCTGAATCAGAAG |
| *chop* reverse | CCACGTGTCTTTTATCTCCC |
| *fgf21* forward | TCGATGATCAAGACAAGCTGA |
| *fgf21* reverse | GCAATTCCTGAAAGGTGCAG |
| *htra2* forward | TGACAGCAGGTATTTCCTTCG |

| | |
|---|---|
| *htra2* reverse | TGTAACGCCTTTTCGATCCT |
| *clpp* forward | AGCAACAATAAGCCGATTCAC |
| *clpp* reverse | ATAGATGGCAGGTCCAGACG |
| *cox4i2* forward | CAAGAGCCTGAAGCAGAAGG |
| *cox4i2* reverse | GGTTTCTTCATCTCCGCAA |
